# Spatiotemporal Dynamics of Single-stranded DNA Intermediates in *Escherichia coli*

**DOI:** 10.1101/2023.05.08.539320

**Authors:** Megan E. Cherry, Katarzyna Dubiel, Camille Henry, Elizabeth A. Wood, Sarah A. Revitt-Mills, James L. Keck, Michael M. Cox, Antoine M. van Oijen, Harshad Ghodke, Andrew Robinson

**Author notes:** To whom correspondence should be addressed. Mailing address: School of Chemistry and Molecular Bioscience, University of Wollongong, Wollongong, NSW 2522, Australia. Correspondence may also be addressed to.

## Abstract

Single-stranded DNA gaps form within the *E. coli* chromosome during replication, repair and recombination. However, information about the extent of ssDNA creation in the genome is limited. To complement a recent whole-genome sequencing study revealing ssDNA gap genomic distribution, size, and frequency, we used fluorescence microscopy to monitor the spatiotemporal dynamics of single-stranded DNA within live *E. coli* cells. The ssDNA was marked by a functional fluorescent protein fusion of the SSB protein that replaces the wild type SSB. During log-phase growth the SSB fusion produces a mixture of punctate foci and diffuse fluorescence spread throughout the cytosol. Many foci are clustered. Fluorescent markers of DNA polymerase III frequently co-localize with SSB foci, often localizing to the outer edge of the large SSB features. Novel SSB-enriched features form and resolve regularly during normal growth. UV irradiation induces a rapid increase in SSB foci intensity and produces large features composed of multiple partially overlapping foci. The results provide a critical baseline for further exploration of ssDNA generation during DNA metabolism. Alterations in the patterns seen in a mutant lacking RecB function tentatively suggest associations of particular SSB features with the repair of double strand breaks and post-replication gaps.

## INTRODUCTION

Single-stranded DNA (ssDNA) is formed as an intermediate during many types of DNA replication and repair transactions. During replication, ssDNA intermediates include lagging-strand gaps [1, 2] and lesion-containing post-replication gaps formed via lesion-skipping [3-5]. Short regions of ssDNA in the form of transcription bubbles are common [6-9]. DNA repair processes such as mismatch repair [10, 11], base excision repair [12], nucleotide excision repair [13, 14], and double-strand break repair [15, 16] all generate ssDNA intermediates of varying length. Few techniques exist for detecting these ssDNA intermediates in live cells [17]. As a consequence, we currently have limited understanding of how ssDNA-producing pathways interact with each other and with other processes that take place on DNA. That limited understanding extends to the frequency of many types of events that generate ssDNA during normal growth as well as the lengths of ssDNA gaps generated by some processes.

A more systematic exploration of the ssDNA genomic landscape was recently initiated using a new whole-genome sequencing approach based on non-denaturing bisulfite treatment followed by deep sequencing [17]. This enabled the first quantitative measurements of ssDNA regions in *E. coli* cells. By this approach it was determined that regions of ssDNA form frequently during normal log-phase growth. These regions are equally distributed between the Watson and Crick strands, and between the leading and lagging strands. Unsurprisingly, most ssDNA gaps are associated with replication as evidenced by a monotonic gradient of ssDNA gap frequency extending from the origin to the terminus of replication [17]. Irradiation of cells with UV light increased the number of ssDNA-containing regions and the lengths of ssDNA tracts.

As part of our broader inquiry into the genomic landscape of ssDNA, we now describe a second method that complements the population-averaged sequencing data with single-cell microscopy data, obtaining information on the localization and dynamics of ssDNA within cells. We used a fluorescent-protein fusion of the ssDNA-binding protein SSB to directly visualize regions of ssDNA [18]. *E. coli* SSB is a stable homo-tetramer composed of 177 amino acid monomers [19]. As a key protein in the initial steps of ssDNA processing, SSB is a clear candidate for use as an *in vivo* gap probe. SSB is essential for genome stability, coating regions of ssDNA to prevent nucleolytic attack and aberrant intra-strand pairing [20-23]. Each SSB monomer contains an N-terminal domain bridged to a C-terminal tail by an intrinsically disordered linker (IDL) region [19, 24]. The 112 amino acid N-terminal domain enables tetramerization and contains the OB fold with which ssDNA interacts [24]. The highly conserved C-terminal tail functions as an interaction site for a multitude of binding partners [21, 25]. These SSB-protein interactions function to recruit and sometimes stimulate DNA replication and repair proteins [25-33]. The unstructured IDL region, which plays a part in SSB cooperativity and DNA binding, is not essential for all SSB functions [19, 24, 34-37]. A variant in which much of the IDL region is removed supports normal growth in *E. coli* [35, 38].

The Keck lab recently developed novel labelled SSB constructs in which the fluorescent proteins mTur2 and sfGFP were individually spliced into the IDL region of SSB [18]. The fusions exhibit binding to short ssDNA oligonucleotides as well as to the SSB-interacting protein ExoI that is similar to wild-type SSB. Additionally, the fusion constructs are functional in a single-molecule rolling-circle replication assay and for strand-displacement synthesis by Pol III. Cells expressing SSB-mTur2 or SSB-sfGFP in place of wild-type SSB exhibit similar growth rates, protein stability, and fitness as the wild-type parent. They exhibited only a modest DNA repair defect [18]. The goal of the current work is to establish a detailed baseline for the use of this SSB variant to explore genomic ssDNA dynamics.

In this study we used the SSB-mTur2 fusion to directly visualize ssDNA gaps in *E. coli* cells. SSB-bound tracts of ssDNA will present as foci in microscopy images. We expect that ssDNA gaps less than the minimal ssDNA binding site for SSB (~35 nt in length [24, 25, 39, 40]) will not attract substantial SSB binding. The ssDNA tracts produced at transcription bubbles (16-18 nt; [6-9]), as well as the intermediates of base-excision repair (5-6 nt; [12]) and nucleotide-excision repair (12-13 nt; [13, 14]) are short and transient enough to limit SSB binding. Lagging-strand gaps (up to 2000 nt; [1, 2]) should attract sufficient SSB to present as foci in microscopy images and render replisomes reliably visible. Similarly, lesion-containing post-replication gaps generated by lesion-skipping [3-5] should be long enough to attract SSB binding. Although detailed information about their lengths is not yet available, observations made by Romero et al. suggest a minimum average gap size of 300 nt for these species [41]. The often long ssDNA gaps produced during mismatch repair [10, 11] should also yield SSB foci. The ssDNA produced during double-strand break repair can be many thousands of nucleotides long [15, 16] and is expected to be coated with a mixture of SSB and RecA.

Fluorescence microscopy measurements revealed a series of novel SSB features in cells that exhibit complex dynamics on both the low-minutes and hours timescales. Here, we present an extensive characterization of these SSB features, their dynamics, and their positioning relative to replication forks. A wider variety of SSB-enriched structures are formed in cells exposed to UV light.

## MATERIALS AND METHODS

### Construction of strains and plasmids

A complete list of the strains used in this study appears in Table 1. A complete list of plasmids used appears in Table 2. EAW1169 (*ssb-mtur2*) and EAW1173 (*ssb-sfGFP*) have been described previously [18] and were constructed using λ_RED_ recombination. The dual-color strains MEC189 and MEC191 were created by transducing the *dnaQ-mKate2::kanR* allele into EAW1169 and EAW1173, respectively, using standard P1-transduction protocols [42, 43]. A Δ*recB* derivative of EAW1169 (EAW1219) was created by replacing the *recB* gene with a kanamycin cassette flanked by FRT sites using the Datsenko-Wanner protocol [44]. Plasmid pEAW1197 was constructed by digesting pKD039 (pET21a-SSB-mTur2) [18] with EcoRI and NdeI to produce a small *ssb-mtur2* containing DNA fragment, then ligating this fragment into pBAD/Myc-HisA (Invitrogen) cut with the same enzymes. The pBAD/Myc-HisA used for this purpose was previously modified to replace the NcoI site with an NdeI site. Plasmids were transformed into chemically competent cells of the indicated parent strain (Table 1) using standard chemical transformation protocols.

**Table 1.** List of *Escherichia coli* strains used in this study. ^1^See Table 2 for description of plasmids.

| Strain name | Relevant Genotype | Plasmid<sup>1</sup> | Parent Strain | Reference |
| --- | --- | --- | --- | --- |
| MG1655 | *ssb<sup>+</sup> dnaQ<sup>+</sup> recB<sup>+</sup>* | – | – | [68] |
| EAW1169 | *ssb-mtur2::FRT* | – | MG1655 | [18] |
| EAW1173 | *ssb-sfGFP::FRT* | – | MG1655 | [18] |
| EAW1219 | *ssb-mtur2::FRT ΔrecB::kan<sup>R</sup>* | – | EAW1169 | this study |
| MEC189 | *ssb-mtur2::FRT*<br>*dnaQ-mKate2::kan<sup>R</sup>* | – | EAW1169 | this study |
| MEC191 | *ssb-sfGFP::FRT*<br>*dnaQ-mKate2::kan<sup>R</sup>* | – | EAW1173 | this study |
| MEC223 | *ssb<sup>+</sup>* | pEAW1197 | MG1655 | this study |

**Table 2.** List of plasmids used in the study.

| Plasmid name | Description | Parent plasmid | Reference |
| --- | --- | --- | --- |
| pKD039 | *ssb-mtur2* | pET21a | [68] |
| pEAW1197 | *ssb-mtur2* | pBAD/Myc-HisA | this study |

### Plate-reader calibration

A calibration curve was generated to correct raw absorbance readings collected outside the linear range of the plate-reader (POLARstar Omega, BMG Labtech, Germany) used in growth assays, adapting a previously published correction procedure [45, 46]. The absorbance of several sample serial dilution series of MG1655 derived *E. coli* strains was measured against media blanks. Blanked OD_600_ readings (OD_obs_) were plotted against the known dilution factors. Values judged, by eye, to be within the linear range of the equipment were linearly fit (Excel, Microsoft, USA). Corrected OD values (OD_corr_) were generated using this best-fit line. OD_corr_ and OD_obs_ values from a minimum of 3 independent experiments were plotted against each other and fit to a cubic polynomial (Origin 2018, Origin Lab, USA) to yield an equipment specific calibration curve.

### Growth assay

The growth of *E. coli* strains MG1655, EAW1169, EAW1173, MEC189, and MEC191 in rich media was assessed using a plate-reader assay. Strains were cultured in EZ-rich defined media supplemented with 0.2% (w/v) glucose at 37°C. Saturated overnight cultures were reset (1:800) in freshly filtered medium. 200 µL aliquots of the resulting cultures were then plated in technical triplicate against a cell-free blank in ultra-low attachment 96-well plates (CL3474, Corning, USA). An oxygen permeable membrane (BEM-1, Diversified 581 Biotech, USA) was applied to ensure sterility and minimize evaporation over time. Plates were then incubated in the plate-reader (POLARstar Omega, BMG Labtech, Germany) for 16 h. Absorbance at 600 nm (OD_600_) was measured every 20 min with bi-orbital shaking (200 rpm) 60 s prior to each measurement. A minimum of 3 independent biological replicates, in technical triplicate, were conducted for each strain.

Before generating growth curves, it was first necessary to correct raw absorbance measurements. Raw absorbance measurements were corrected by subtracting the mean blank absorbance at each time point from the corresponding mean OD_600_ of each technical triplicate set (OD_obs_). A plate-reader specific calibration curve was then applied yielding the corresponding OD_corr_ values. OD_corr_ values were then normalized between 0 and 1 across all time points (OD_norm_). To generate growth curves, OD_norm_ values were fit to a 4-parameter Richard’s curve (Equation 1) in Origin 2018 (Origin Lab, USA). In this model, *a* is defined as the top asymptote, *xc* as the x-axis inflection-point, *d* as an inflection coefficient, and *k* as the growth rate. As wild-type MG1655 cells cultured in EZ-rich defined media grew to saturation in less than 16 h, only OD_norm_ values from the first 6 h were included as the fitting data set.

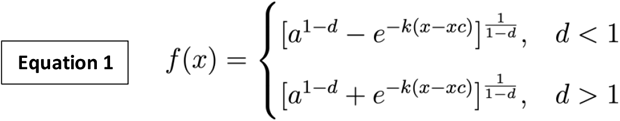

### UV survival assay

The survival of *E. coli* strains MG1655, EAW1169, EAW1173, MEC189, and MEC191 following UV exposure was assessed using a spot plate assay adapted from a previously published protocol [47]. For each strain, a streak purified colony was grown, in 1 mL of LB in 2 mL microcentrifuge tubes, at 37 °C between 16 – 24 h at 800 rpm (ThermoMixer C, Eppendorf, Germany). 10 µL of saturated culture was then reset in 1 mL of LB before incubating an additional 2 h. Exponential phase cells (OD_600_ = 0.5-0.6) were then pelleted at 5000 rpm (Centrifuge 5425, Eppendorf, Germany) for 5 min at room temperature. Cell pellets were washed twice with PBS before resuspending in PBS to a final OD_600_ of 0.25. 30 µL (∼6 × 10^6^ cells) droplets of each strain were placed onto a sterilized petri dish lid and irradiated (Herolab UV-8 SL, Herolab, Germany) with 254 nm UV light at doses of 0, 3, 10, and 30 J/m^2^. 10-fold serial dilutions of irradiated cells were made in PBS. 5 µL spots of these serial dilutions (10^−1^ to 10^−8^) were then incubated on LB Agar plates at 37 °C for 16 h. A minimum of two biological replicates were conducted for each strain and irradiation condition. UV flux was measured using a UVX radiometer (UVP, USA) prior to irradiation in all experiments.

### Preparation of APTES-functionalized microscope coverslips

APTES (3−Aminopropyltriethoxysilane) functionalized slides were prepared as previously described [48]. To clean, glass slides (No. 1, 24 mm × 55 mm, 0101232, Marienfeld, Germany) were sonicated in 20 – 40% (w/v) KOH for 20 min at room temperature and rinsed extensively with fresh Milli-Q water. After cleaning, coverslips were incubated in freshly prepared 5% (w/v) APTES for 12 – 20 min at room temperature. Slides were then sonicated in ethanol for 10 – 30 s prior to rinsing with Milli-Q water. Nitrogen-dried functionalized coverslips were stored in an airtight container at room temperature for a maximum of 24 h prior to use.

### Flow cell assembly

Live-cell fluorescence imaging was performed in custom-built flow cells, prepared as previously described [48]. Briefly, inlet tubing (BTPE-60, Walker Scientific, AUS), cut to 30–45 cm, was glued (BONDiTTM B-482 epoxy, Reltek, USA) to quartz slides and trimmed flush once dry. Scotch double-sided adhesive transfer tape (970XL, 3M, USA), arranged in parallel strips along the long edges of the quartz top, was used to attach the quartz top to an APTES-functionalized coverslip. The resulting closed channel was further sealed with 5-minute epoxy (Parfix, AUS) applied along all exposed quartz/glass and quartz/tubing interfaces. Once dry, a straight-edge razor was used to remove excess glue from the bottom of the flow cell. Custom built flow cells were stored on the bench-top at room temperature for a maximum of 24 h prior to use in imaging experiments.

### Cell culture for imaging

For all imaging experiments, cells were grown at 37 °C for 90–120 min to exponential phase (OD600 = 0.5 – 0.6) in EZ-rich defined medium. Cells carrying pEAW1197 were grown in the presence of ampicillin (100 µg/mL) in media supplemented with 0.2% (w/v) glycerol. Expression of *ssb-mtur2* in pEAW1197 containing strains was induced 30 min prior to imaging by the addition of 0.002% L-arabinose. All other cells were grown in media supplemented with 0.2% (w/v) glucose.

Cells were manually loaded into quartz-top flow cells at 37 °C by flowing in cell culture and allowed several minutes to adhere on the APTES functionalized surface in the absence of flow. Following this, a syringe pump (Adelab Scientific, Australia) was used to deliver a constant supply of freshly aerated growth media to the flow cell to both dislodge loosely associated cells and to supply a flow of fresh medium. In experiments utilizing *in situ* UV irradiation, UV light was delivered at a fluence of 10 J/m^2^ via a mercury lamp (λ = 254 nm, UVP).

### Fluorescence microscope and imaging

Single-molecule fluorescence imaging measurements were performed on a custom wide-field microscope comprised of a Nikon Eclipse Ti-2 body, a 1.49 NA 100x objective, 458-nm (75 mW max. output) and 568-nm (200 mW max. output) Sapphire LP lasers (Coherent, USA) and a 512 × 512 pixel^2^ EM-CCD camera (C9100-13, Hamamatsu, Japan).

Rapid acquisition movies (600 × 50 ms frames, continuous excitation with 458 nm light; power density: 150 W/cm^2^) were collected to determine the number of SSB-mTur2 molecules per cell. Emission from mTur2 was collected between 468 and 495 nm (ET 485/30m filter, Chroma). Time-lapse movies were recorded to characterize changes in cellular SSB-mTur2 levels and to measure colocalization of these molecules with the replisome marker, DnaQ-mKate2. mKate2 was imaged using a 568 nm laser excitation at a power density of 180 W/cm^2^ and collected between 610 - 680 nm (ET 645/75m filter, Chroma).

Time-lapse image sets with 30 s intervals were recorded in sets of two (brightfield [34 ms exposure], mTur2 fluorescence [100 ms exposure; 15 W/cm^2^]) over a course of 15 min. For these relatively rapid time-lapse sets, fluorescence imaging conditions were chosen to minimize photodamage. In instances where cells were UV irradiated, cells were dosed immediately after the acquisition of an initial pre-UV round of images.

Time-lapse image sets with 10 min intervals were recorded in sets of three (brightfield [34 ms exposure], mTur2 fluorescence [500 ms exposure; 150 W/cm^2^], mKate2 fluorescence [200 ms exposure; 180 W/cm^2^]) over a course of 2 h. For these relatively slow time-lapse sets, fluorescence imaging conditions were chosen to maximize signal-to-noise. In instances where cells were UV irradiated, cells were dosed immediately after the acquisition of an initial pre-UV round of images. Additional time-lapse movies were recorded to characterize changes in mTur2 fluorescence in the flow cell environment over time in the absence of exogenous damage. In these experiments, mTur2 fluorescence [500 ms exposure] was collected over 5 × 100 ms frames.

### Image processing

Image processing was performed in Fiji v1.51n [49], using the Single-Molecule Biophysics plug-in (available at https://github.com/SingleMolecule/smb-plugins) and custom macros. Custom Fiji macro *Process Raw Images* was used to convert raw ND2 images to TIF format. Background correction and image flattening was then performed as previously described [50-52]. These corrections were necessary due to excitation inhomogeneity and extraneous background signal originating from the flow cell surface. To address excitation inhomogeneity, for each experiment a normalized beam profile was created from the average projection of the concatenated stack of all images taken over all time points. As needed for higher intensity signal, a median filter (radius = 100 pixels) was applied for further smoothening prior to the normalization of beam profile intensities. Individual images were then divided by the normalized beam profile image to flatten intensities across the image. Once flattened, flow cell auto-fluorescence, estimated as the median intensity on each frame, was subtracted prior to remapping all intensities across 16-bits. For ease of processing, the Grid/Collection stitching [53] Fiji plug-in was used to stitch together flattened image stacks across multiple positions.

MicrobeTracker 0.937 [54] was used to manually create cell outlines as regions of interest (ROIs). Manual outlines were used to ensure that only non-overlapping, in-focus cells were selected for analysis. Cell metrics such as cell length, area, and volume, were also generated using this plug-in. Cell outline ROIs were imported into Fiji to extract additional cell metrics including mean cell intensities and foci per cell.

### Analysis of mTur2 single-molecule integrated density

The mTur2 single-molecule intensity was extracted by applying change point analysis (34) to SSB-mTur2 focus photo-bleaching trajectories. Trajectories of long-lived SSB binding events were generated in Fiji v1.51n [49] using the *Single-Molecule Biophysics* plug-in and custom macros. Long-lived binding events were identified by taking the average projection, across all frames, of the discoidal averaging filtered flattened image stack (inner radius of one pixel, outer radius of four pixels). *Peak Fitter* was then used to detect foci in this projected image above a specified threshold. Custom Fiji macro *Changepoint Peaks* was then used to extract mean intensity photo-bleaching trajectories at each peak location in the original flattened rapid acquisition stack.

Any raw intensity measurement of cell features is the linear combination of cellular auto-fluorescence, flow cell surface background signal, and the desired fluorophore fluorescence signal. To ascertain the desired mTur2 contribution, extracted mean intensities were calculated as the difference between the mean intensities measured in rectangular ROIs centered at the peak locations with inner and outer radii of 2 and 4 respectively.

To distinguish discrete photo-bleaching steps from noise, a change-point algorithm (34) was applied to each peak photo-bleaching trajectory (sigma value = 15). Single-molecule step size (I_step_) histograms were then built in MATLAB (MathWorks, USA), using only the final intensity step from each mean intensity photo-bleaching trajectory. The mean value for I_step_ was determined by a single term Gaussian fit to the binned data. The mTur2 single-molecule integrated density was then calculated as the product of the mean pixel I_step_ and the pixel area of a diffraction limited spot (16 px^2^).

### Analysis of SSB cell copy numbers

Copy numbers of SSB were calculated by dividing the background-free integrated density of chromosomally expressed SSB-mTur2 by the mTur2 single-molecule integrated density. To extract background-free integrated densities of SSB-mTur2 in cells, it was first necessary to calculate the signal contribution of cellular auto-fluorescence. A custom Fiji macro (*MeasureROIsAllFrames*) was used to generate mean intensity trajectories for MG1655 cells. Intensities were measured in flattened image stacks inside manual cell outlines exported from MicrobeTracker 0.937 [54]. OriginPro 2018 was then used to globally fit each intensity trajectory to a single exponential function, **Equation 2**.

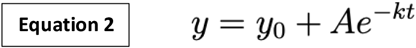

For each cell, *k* is the photo-bleaching rate and amplitude *A* is an accurate measure of the mean auto-fluorescence intensity per pixel. As *k* and *A* were similar across all imaging growth conditions, the mean auto-fluorescence pixel intensity was determined as the mean *A* of the combined data set.

Raw integrated densities of SSB-mTur2 signal were measured from the first image in each MEC189 (*ssb-mtur2 dnaQ-mK2*) flattened rapid acquisition image stack. The background-free integrated density was then calculated as difference of the raw integrated density and the auto-fluorescence integrated density (auto-fluorescence pixel intensity × cell area). For each cell, the SSB-mTur2 copy count was then calculated by dividing the corrected integrated density by the mean mTur2 single-molecule integrated density. The cellular concentration was calculated using the volume of each cell, determined from cell outline assignment in MicrobeTracker [54].

### Characterization of SSB features

Custom Fiji macro, *Particle Analysis Suite v4* was used to detect and characterize SSB cellular fluorescence and features. Cell outline regions of interest (ROIs), exported from MicrobeTracker 0.937 [54], were used to measure mean cell fluorescence intensities. Bound SSB features were categorized as either isolated diffraction-limited peaks or clusters of indistinguishably grouped peaks, heretofore referred to as clusters.

SSB cluster ROIs were identified by thresholding as described below. First, fluorescence not attributable to in-focus cells was removed from flattened image stacks. To do this, cell outlines were used to generate positive binary masks of all stitched image stacks. In these masks, area contained within cell outlines was assigned an intensity of 1 with all other areas having intensity of 0. Fiji Image Calculator was then used to apply these binary masks to the corresponding flattened stitched image stacks, eliminating any intensity not contained within cell outlines. Next, diffuse SSB intensity was filtered out with a discoidal filter of inner radius 1 pixel and outer radius 8 pixels. Software limitations required the conversion of 16-bit stacks to 8-bit before thresholding intensities. Thus, after arbitrarily setting the 16-bit intensity range to 0– 1500, processed image stacks were converted to 8-bit and thresholded to identify high intensity regions. An 8-bit intensity threshold of 175 -255 utilizing Fiji’s default thresholding algorithm, was found to best capture bound SSB particles. Lastly, cluster ROIs were generated using the Analyze Particles dialogue for thresholded regions with area of >16 px^2^. Cluster characteristics, including area, mean fluorescence intensity, centroid, and circularity were extracted in Fiji [49]. Centroid positions were then used to assign clusters to cells in MicrobeTracker [54].

SSB peak foci were detected using Peak Fitter from the *Single-Molecule Biophysics* Fiji plugin. Briefly, a discoidal averaging filter of inner radius 1 pixel and outer radius 8 pixels was first applied to stitched flattened image stacks to filter out diffuse signal. A Gaussian was then fit to pixel groupings above a set intensity threshold (mean + 6x standard deviation), with a fit radius of 3 pixels and assumed minimum distance of 4 pixels between peaks. 4 × 4 pixel peak ROIs, centered on each peak location, were then generated for each peak location. Only peak locations contained within cell ROIs, but excluded from cluster ROIs, were used for analysis.

DnaQ-mKate2 foci were detected using Peak Fitter from the *Single-Molecule Biophysics* Fiji plug-in. A discoidal averaging filter of inner radius 1 pixel and outer radius 4 pixels was applied before fitting a Gaussian to pixel groupings above a set intensity threshold (250), with a fit radius of 4 pixels and minimum distance of 4 pixels between peaks. 4 × 4 pixel peak ROIs, centered on each peak location, were then generated for each peak location. Only peak locations contained within cell ROIs were used for analysis.

### Analysis of SSB/replisome overlap

Custom Fiji macro, *Particle Analysis Suite v4* was used to calculate overlap statistics between SSB and replisome foci. Two features were classified as overlapping if any portion of both ROIs occupied the same space within a cell. The overlap frequency for SSB with replisomes and replisomes with SSB was then reported as the respective number of features (either SSB or replisomes) with overlapping area divided by the total number of features. A custom MATLAB (MathWorks, USA) script [47] was used to perform frequency calculations two hundred times on data sets derived from randomly selecting, without replacement, 80% of events. The percentage of overlapped area and centroid-centroid separation was then calculated for any overlapped features. Though the percentage of overlapped area may be attributable to multiple overlap events, only the minimum centroid-centroid separation distance was reported. Chance colocalization was calculated as the average cell area occupied by either replisome foci or SSB peak and particle ROIs.

## RESULTS

To visualize time-dependent changes in SSB expression and cellular localization, we examined *E. coli* strains expressing a functional fluorescent protein fusion of the SSB protein (SSB-mTur2) as the sole source of SSB in the cell **[18]**. In this construct, the fluorescent protein mTurquoise2 is inserted between amino acids F148 and S149 of the SSB protein, partway through the intrinsically disordered linker (IDL) region. A dual-color strain (MEC189) carrying an additional marker for DNA polymerase III holoenzyme (pol III HE) was also produced. In this strain the red fluorescent protein mKate2 is fused to the C-terminus of the DnaQ protein (ε subunit of pol III HE). This DnaQ-mKate2 fusion has previously been used to detect the intracellular localization of replisomes and has been shown to have no impact on the morphology or growth rate of cells **[18]**. An SSB-sfGFP fusion was also analyzed and returned similar results to the SSB-mTur2 fusion (see Supplementary Material).

### SSB-mTur2 remains functional when cells are grown in imaging medium

Previous characterization of the SSB-mTur2 fusion indicated that cells expressing the fusion exhibited wild type growth rates and fitness in LB medium [18]. Before we started fluorescence microscopy measurements, we characterized the phenotypes of strains growing in the specific medium to be used in the fluorescence microscopy measurements: EZ rich defined medium supplemented with 0.2% glucose. Cell growth parameters were found to be similar for SSB-mTur2 strains and wild-type cells (**Supplementary figure 1A**). Bright-field measurements revealed very slight differences in cell lengths between unlabeled and labelled SSB strains when grown to early exponential phase in EZ-rich defined media supplemented with 0.2% glucose (**Supplementary figure 1B**). These differences were deemed to be negligible in the context of the current study. A spot plate assay was used to assess UV sensitivity at 0, 10, and 30 J/m^2^ (**Supplementary figure 2**). At both 0 and 10 J/m^2^ the wild-type and SSB-mTur2 strains displayed similar survival profiles. Under the conditions of these experiments, most cells (>99%) ultimately survive this dose of UV [55] and this is the dose we applied in the current experiments. At the higher 30 J/m^2^ dose, SSB-mTur2 strains displayed a slight (less than one order of magnitude) UV sensitivity in comparison to wild-type cells. This sensitivity is consistent with observations made by Dubiel *et al*. for cells grown in LB medium [18]. The slight UV sensitivity associated with the SSB-mTur2 fusion was not exacerbated by the presence of the DnaQ-mKate2 fusion (**Supplementary figure 2**). Overall, results indicate that the SSB mTur2 fusion maintains near-wild type activity in EZ medium, displaying only a slight sensitivity phenotype in response to high doses of UV light. Similar results were obtained for strains expressing an SSB-sfGFP fusion (**Supplementary figure 2**).

### The intracellular concentration of SSB-mTur2 is in the low micromolar range

Using a single-molecule fluorescence microscope it is possible to calibrate the fluorescence intensities of cells to the fluorescence of individual fluorophore molecules. This enables measurement of protein copy numbers and, when used in conjunction with cell morphology measurements, intracellular protein concentrations. The SSB-mTur2 fusion is expressed from the native *ssb* promoter. Western blots presented by Dubiel *et al*. confirm the approximately 3.6-fold higher expression of the fluorescent IDL fusions as compared to EcSSB [18]. Quantifying the intracellular levels of SSB-mTur2 should therefore provide an indication, albeit elevated, of the expression levels of SSB in wild-type cells. In this type of measurement, the value for the mean intensity of a single fluorescent protein is extracted from step-wise photobleaching measurements [50, 51, 56]. We found the SSB-mTur2 protein to be remarkably photostable, bleaching slowly and without producing steps suitable for intensity measurements. As an alternative approach we used an arabinose-inducible plasmid to express very low amounts of SSB-mTur2 within otherwise wild-type cells (**Supplementary figure 3A–E**). This approach provided suitable photobleaching steps and enabled the mean intensity of a single SSB-mTur2 molecule to be determined. Comparing this value to the integrated fluorescence intensities of cells expressing only SSB-mTur2, and accounting for spurious signals arising from cellular autofluorescence (**Supplementary figure 3F–G**), we determined the mean number of SSB-mTur2 molecules per cell to be 3282 monomers (95% CI: 2896 – 3667, n = 72 cells), equating to ~820 tetramers. Accounting for cell volumes, we determined the mean intracellular concentration of SSB-mTur2 to be 1.6 µM (95% CI: 1.4 – 1.7 µM, n = 72 cells). This is about 40% of the level previously estimated from proteome analysis albeit under somewhat different growth conditions [57].

### SSB dynamics under rapid-growth conditions

We monitored SSB-mTur2 signals in cells by time-lapse microscopy, recording images at 30s intervals for 5 min. To minimize photodamage we used short exposures (100 ms) and relatively low laser power (15 W/cm^2^). We started by monitoring cells growing in rich medium in a flow cell at 37 °C, without any exogenous DNA damage. The doubling time under these conditions is 19 min. Cells growing in LB medium have a similar doubling time and previous work suggests they contain an average of 14 replisomes per cell[58]. As the typical number of replisome foci is substantially less than 14, we presume multiple replisomes are often clustered. Under these conditions we primarily expect to observe SSB foci forming at replisomes, and potentially at post-replication gaps and intermediates produced during mismatch-repair and double strand break repair. Regions of ssDNA within transcription bubbles and intermediates of base-excision repair and nucleotide excision repair are too small to bind significant amounts of SSB. The SSB-mTur2 cells were introduced to a home-built flow cell mounted on a bespoke single-molecule fluorescence microscope, as described previously [50-52, 56]. The cells attached to the lower surface of the flow cell, which was comprised of a positively charged APTES-derivatized glass coverslip.

Under normal growth conditions we observed a mixture of SSB-mTur2 foci and diffuse signal spread throughout the cytosol (**Figure 1A–B**). Cells contained 3.0 foci on average (95% CI: 2.3 – 3.7). Commonly, there were two foci or clusters of foci, one near each end of a cell. In most cases, 2-3 foci could be discerned within a cluster. Occasionally foci were produced that were much brighter than the others (**Figures 1A–B**; magenta arrows). Manual inspection of the timelapse videos suggested that the bright foci were formed in two ways: i) multiple foci in a cluster *merged* to form a single, brighter focus (**Figure 1A–B**; arrows labelled *m*); ii) a relatively dull focus *brightened* (**Figure 1B**; arrows labelled *b*). Bright foci typically lasted only a few frames (30–120s) before resolving in one of two ways: i) by *splitting* into a cluster of multiple foci (**Figure 1A–B**; arrows labelled *s*); or ii) by *dulling* (**Figure 1B**; arrows labelled *d*). These behaviors are quantified in the following section.

**Figure 1.**
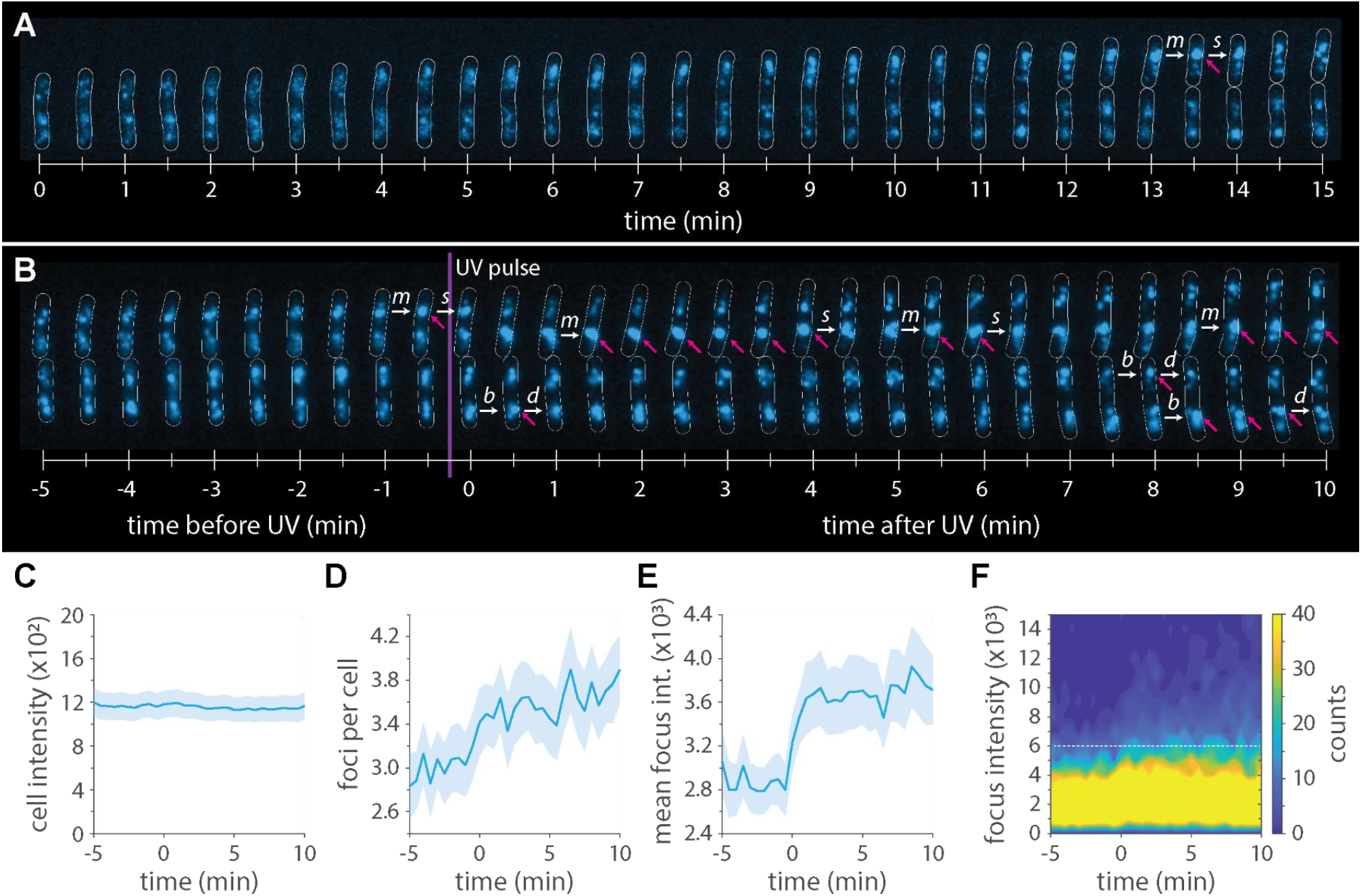
Spatiotemporal dynamics of SSB-mTur2 on the low minutes timescale. *E. coli* cells expressing SSB-mTur2 (EAW1169) were imaged on a single-molecule sensitive fluorescence microscope. (**A–B**) Montages of cells within timelapse series. Magenta arrows indicate ‘bright’ foci, with intensities above 6000 arbitrary units. White arrows indicate dynamics that lead to the production, or dissolution of bright foci: (*b*) brightening of an existing focus; (*m*) merging of multiple foci into a single focus; (*d*) dulling of a bright focus into a regular focus; (*s*) splitting of a bright focus into multiple regular foci. (**A**) Cells growing under normal conditions for 15 min. (**B**) Cells growing under normal conditions for 5 min then for 15 min following irradiation with light (10 J/m2 delivered over 10s). (**C**) Plot of mean intracellular SSB-mTur2 signal intensities versus time. Shaded area indicates 95% confidence intervals. (**D**) Mean number of SSB foci per cell plotted as a function of time. Shaded area indicates 95% confidence intervals. (**E**) Plot of mean intensities of SSB-mTur2 foci versus time. Focus intensities were determined by fitting the intensity profiles of foci with two-dimensional Gaussian functions. The height parameter is plotted here. Shaded area indicates 95% confidence intervals. (**F**) Two-dimensional histogram of focus intensities plotted as a function of time. White dotted line indicates the threshold value used to assign ‘bright’ foci.

### UV-induced SSB dynamics on the low-minutes timescale

We next examined cells after exposing them to a flash of short wavelength UV light (10 J/m^2^; λ = 254 nm) through the UV-transmissible quartz top piece of the flow cell. The timelapse imaging (see section above) was continued for a further 10 minutes after UV irradiation. Under these conditions we anticipated that SSB focus dynamics might change as replisomes began to encounter UV-induced lesions in the template DNA.

The overall intensity of SSB-mTur2 signal within cells did not change upon UV irradiation (**Figure 1C**), indicating that the intracellular concentration remained relatively constant before and after exposure. Analysis of the number of foci per cell (**Figure 1D**) suggested a slight increase in the number of foci after irradiation (from 3.0 to 3.6 foci per cell [95% CI: 2.8 – 4.2]), however this was not deemed statistically significant. More strikingly, there was a decrease in the amount of diffuse SSB-mTur2 signal immediately after UV exposure (**Figure 1B**), and the mean intensities of foci increased dramatically (**Figure 1E**). These observations indicate that a significant portion of cytosolic SSB-mTur2 molecules redistributed to the nucleoid in response to UV irradiation. A two-dimensional histogram of focus intensities plotted as a function of time (**Figure 1F**) indicated that the increase in average focus intensities was driven by the appearance of a population of very bright foci, as opposed to all foci uniformly increasing in intensity. To facilitate further analysis, we sought to assign this new population to a distinct class. Prior to UV irradiation, the vast majority of foci displayed intensities not exceeding 6000 arbitrary units (**Figure 1F**, dotted white line). Thus, all foci below this arbitrary threshold represented foci of types present during normal growth. After irradiation, a significant population of brighter foci appeared that exceeded this brightness threshold, often by a substantial margin. Examples of these “bright” foci are illustrated in **Figure 1A-B** [magenta arrows].

Under this definition, 7.6% (95% CI: 7.4 – 7.8%) of foci were classed as bright before UV irradiation. This increased to 17.1% (95% CI: 16.7 – 17.5%) after UV irradiation. We next examined the events leading to the formation of a bright focus, classifying the formation event as either a *merging* or *brightening* event (see section above; indicated in **Figure 1A–B** by arrows labelled *m* or *b*). Of the bright foci that were formed before UV irradiation, 47% were the result of merging events and 53% arose by brightening. Very similar proportions were measured after UV irradiation with 48% of bright foci forming by merging and 52% resulting from brightening events. We next examined the lifetimes of bright foci by noting when each focus rose above, and fell below, the threshold used to assign bright foci. We compared lifetimes before and after UV treatment, and lifetimes for foci formed by merging and brightening (**Supplementary figure 4**). In all cases the lifetimes followed a similar (approximately exponential) distribution; there were no obvious differences in lifetimes for bright foci appearing before or after UV irradiation, or for bright foci formed via merging or brightening. Overall, the lifetime analysis indicated that the bright foci most commonly persist for <30 s but can occasionally persist for >7 min. Classifying the events that lead to resolution of a bright focus as *splitting* or *dulling* (see section above; indicated in **Figure 1A–B** by arrows labelled *s* or *d*), we observed that 49% of bright foci were resolved by splitting and 51% were resolved by dullening. Bright foci formed by merging were overwhelmingly resolved by splitting (97%) while bright foci formed by brightening were overwhelmingly resolved by dulling (96%).

### Dynamics of long-lived SSB features on longer timescales

We next characterized SSB focus dynamics specifically at replication forks using a strain that expressed both SSB-mTur2 and the replisome marker DnaQ-mKate2. Due to rapid photobleaching of the DnaQ-mKate2 probe, it was not possible to record dual-color timelapse movies with short interval times. We instead turned our attention to longer-term SSB dynamics that occur during recovery from UV exposure, recording timelapse movies of SSB-mTur2 DnaQ-mKate2 cells with 10 min intervals, for a total duration of 120 min after UV exposure. As the longer imaging interval decreased the risk of photodamage to cells, we were able to use a longer exposure time (500 ms) to specifically capture longer-lived SSB binding events. A higher laser power (150 W/cm^2^) was implemented to improve the signal-to-noise ratio within images.

We first characterized cell morphology and SSB concentrations (monitored by cellular mean pixel intensity measurements) in cells irradiated with 10 J/m^2^ UV light. Cellular fluorescence and morphology remained unchanged until 40 min after irradiation at which point marked increases in cell length and cellular mean pixel intensity were observed (**Figure 2A; Supplementary figure 5**). These increases persisted through the end of image acquisition, 120 min after irradiation (**Figure 2A**; **Supplementary figure 5**). By the 120 min timepoint, cell lengths had on average increased nearly three-fold (**Supplementary figure 5**). This response is similar to the response observed for wild-type cells, except that wild-type cells began to decrease in cell length from 100 min after irradiation (**Supplementary figure 5**). Fluorescence levels remained elevated 2 h after irradiation, measuring approximately 1.6-fold higher than pre-UV levels. We had previously measured the intracellular concentration of SSB-mTur2 in untreated cells to be 1.6 µM. Multiplying this value by the 1.6-fold intensity increase returns an approximate value of 2.6 µM for the intracellular SSB-mTur2 concentration 120 min after irradiation.

**Figure 2.**
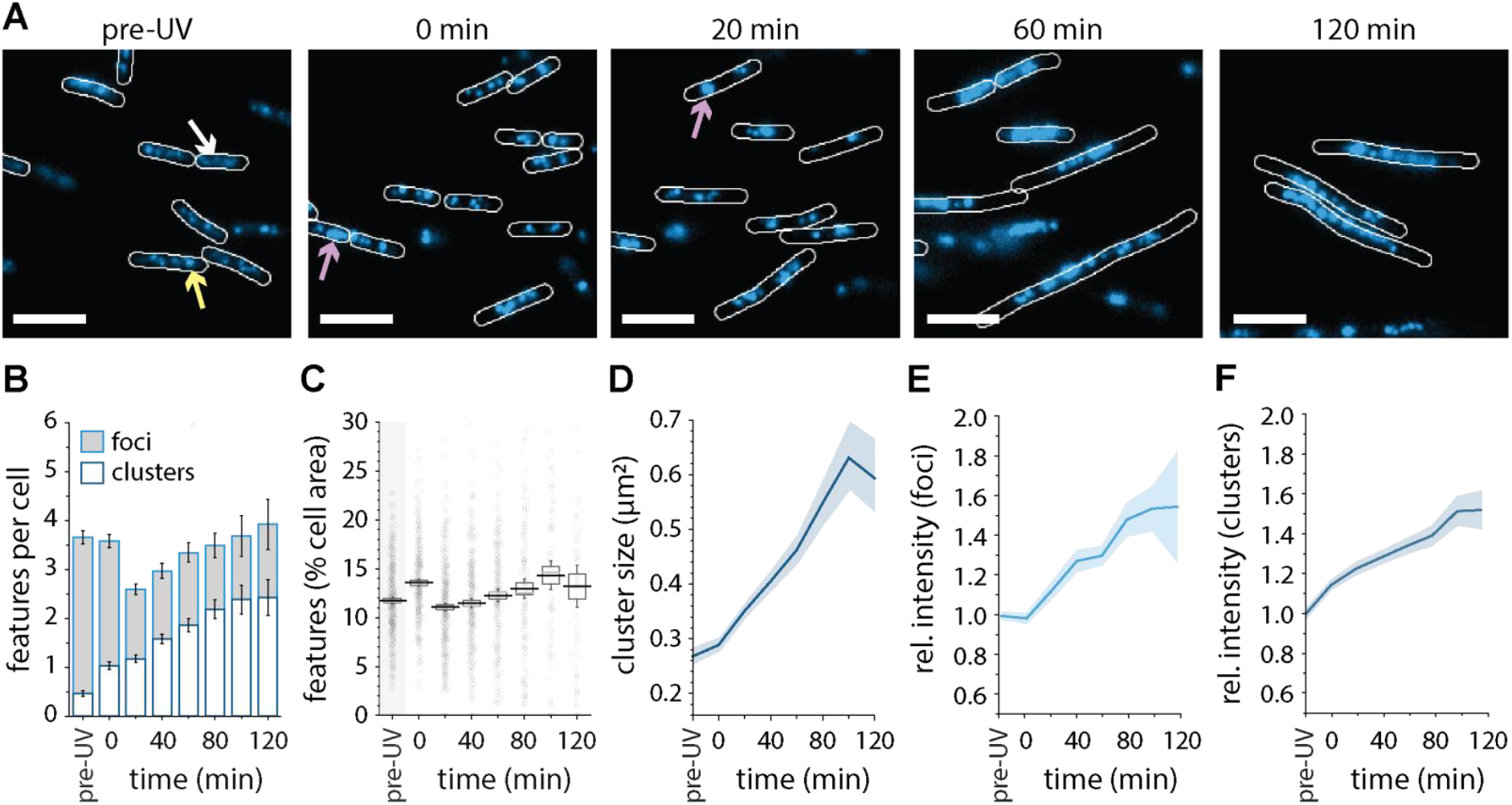
Spatiotemporal dynamics of SSB-mTur2 during recovery from UV *in situ*. **(A)** Example fields of SSB-mTur2 expressing cells. SSB-mTur2 signal was classified into three categories: (i) diffuse, unbound protein (example indicated with white arrow); (ii) diffraction-limited foci (yellow arrow); (iii) large SSB features, (purple arrow). Scale bars indicate 5 µm. (**B**) Proportion cell area occupied by SSB features plotted as a function of time. Plots indicate the mean (black line), median (gray line), 95% CI (whiskers), and SE of the mean (box). (**C**) Quantification of SSB features as a function of time. The distribution of large SSB features (white) and foci (gray) are shown for each time point as a proportion of the total number of SSB features per cell. Error bars indicate 95% confidence intervals. (**D**) Particle size as a function of time. Means indicate the average particle ROI size at the indicated time after UV irradiation. (**E**–**F**) Mean pixel intensities of SSB features as a function of time for foci (**E**) and large SSB features (**F**). Means indicate the average pixel intensity of peaks and particles as fold increases from the pre-UV intensity. Shaded regions indicate 95% confidence interval regions.

We next measured the number and type of SSB features present in cells before and after irradiation with UV light (**Figure 2**). It was clear from the timelapse images (**Figure 2A**) that large SSB features formed in response to UV, most of which appeared to be comprised of multiple partially overlapping foci (**Figure 2A**). We therefore separated SSB features into two classes. From this point in the study, foci were defined as features occupying an area ≤16 pixels^2^ (0.4 × 0.4 µm), the area of a diffraction limited spot. Features larger than this were classified as *clusters*. Prior to irradiation cells contained an average of 3.7 (95% CI: 3.5 – 3.8) SSB features (**Figure 2B**). Of these, 86% were classed as foci and 14% were classed as clusters. The mean intensity of the clusters was, on average, 2-fold higher than that of the foci.

Continued creation and expansion of clusters was observed as cells began to filament, 40 minutes after UV exposure (**Figure 2A**). Though the proportion of cell area occupied by these clusters remained at or near pre-UV percentages until imaging stopped 120 min after irradiation (**Figure 2C**), the total number of clusters present, their sizes, and their intensities all trended upwards until 80 min post-UV (**Figure 2D–F**). For each of these metrics, elevated levels persisted until the end of the imaging window. At 120 min after UV irradiation the total number of features present in each cell were, on average, at or above pre-UV levels (**Figure 2B**); however, the proportion of clusters increased 5-fold. On average the areas of these clusters were more than 2-fold higher than those observed prior to UV irradiation (**Figure 2D**). Furthermore, both foci and clusters displayed increased mean pixel intensities (**Figure 2E–F**), indicating that the density of SSB within each feature was significantly increased.

### Characterization of SSB colocalization with replication forks

Next, the dual-color SSB-mTur2 DnaQ-mKate2 strain was used to assess both the prevalence and localization of DnaQ-mKate2 both before and after cells were irradiated with UV light (**Figure 3; Supplementary figure 6**). From the timelapse images it appeared that: i) there were fewer DnaQ foci than SSB features at all time points, and ii) many of the DnaQ foci localized to the outer edge of SSB clusters (**Figure 3A**). Quantitative analyses were carried out to test these ideas.

**Figure 3.**
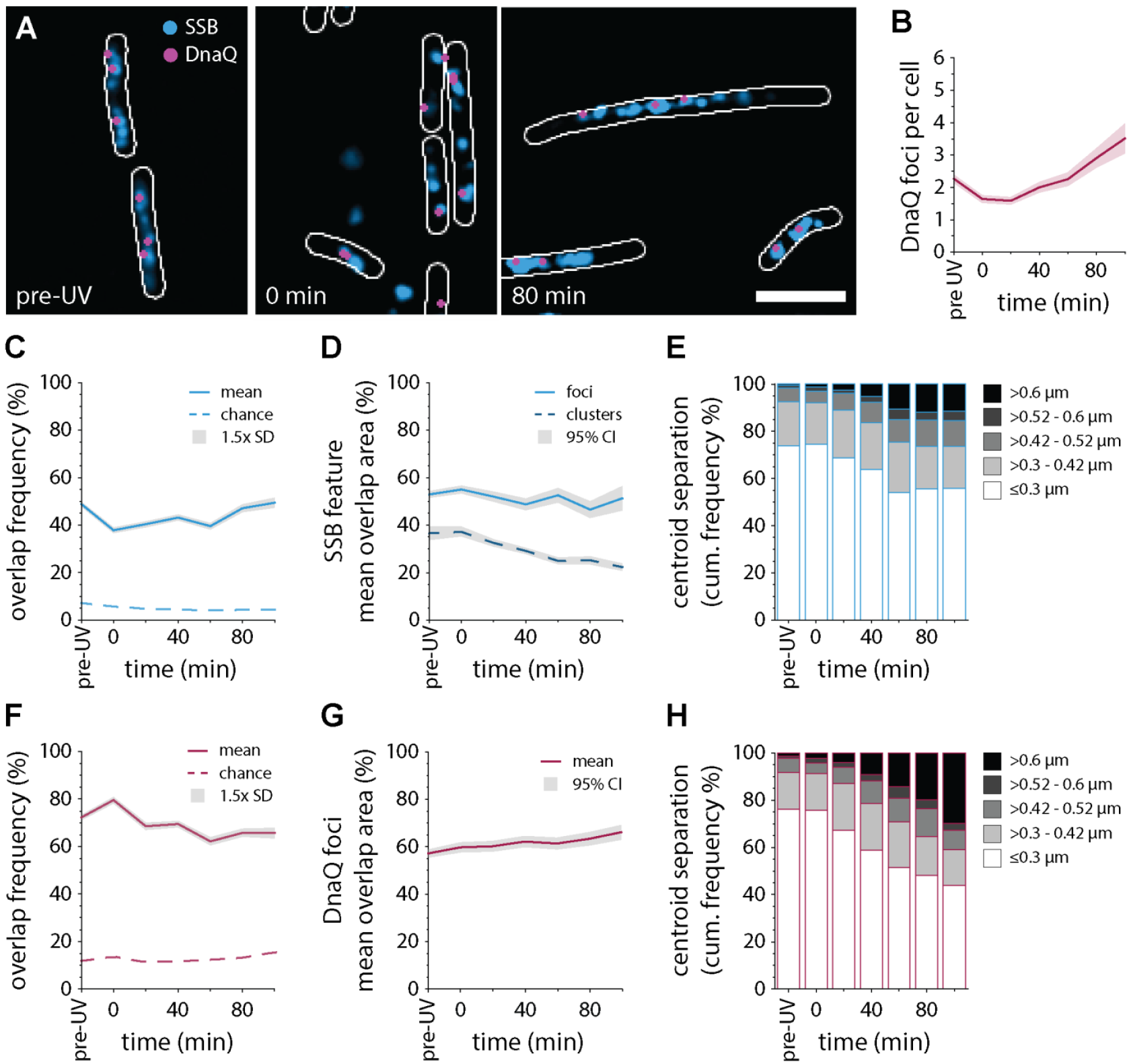
Localization of DnaQ-mKate2 at outer edge of SSB features. **(A)** Example two-color (SSB-mTur2, blue; DnaQ-mKate2, magenta) images recorded prior to, immediately after, and 80 min after irradiating cells with UV light. Scale bar indicates 5 µm. (**B**) Number of DnaQ-mKate2 foci per cell plotted as a function of time after UV irradiation. Shaded region indicates values within 1.5× SD of the mean. (**C**) The percentage of SSB-mTur2 features that spatially overlap with a DnaQ-mKate2 focus, plotted as a function of time. Shaded regions indicate 95% confidence intervals. (**D**) Percent area of each colocalized SSB-mTur2 feature that is overlapped by a DnaQ-mKate2 focus. (**E**) Distances between the centroids of SSB-mTur2 features and the centroids of colocalized DnaQ-mKate2 foci. (**F**) The percentage of DnaQ-mKate2 foci that spatially overlap with an SSB-mTur2 feature, plotted as a function of time. Shaded regions indicate 95% confidence intervals. (**G**) Percent area of each colocalized DnaQ-mKate2 focus that is overlapped by an SSB feature. (**H**) Distances between the centroids of DnaQ-mKate2 foci and the centroids of colocalized SSB features.

Prior to irradiation cells contained an average of 2.3 (95% CI: 2.1 - 2.4) DnaQ foci (**Figure 3B**), compared with 3.7 (95% CI: 3.5 – 3.8) SSB features (**Figure 2B**). Immediately after UV irradiation, a decrease in the number of DnaQ foci was observed (**Figure 3B**). This drop in observed foci coincided with significant decreases in both the proportional cell area occupied by these foci as well as the overall cellular mean pixel intensity associated with DnaQ-mKate2 signal (**Supplementary figure 7**). These observations are similar to observations made previously in the context of ciprofloxacin-mediated DNA-damage (36) and are consistent with UV irradiation promoting the dissociation of some replisomes. After 20 min, the number of observed replisomes remained low, with cells containing 1.6 (95% CI: 1.5 - 1.8) DnaQ foci. Though steady increases in the number of DnaQ foci were observed over time, the density of these foci remained unchanged 100 min post-UV (Supp). Modest increases in the cellular mean pixel intensity were also observed (**Supplementary figure 7**). Signal associated with mKate2 fluorescence returned to pre-UV levels 60 min post-UV.

We next analyzed time-dependent changes in spatial overlap between the observed SSB features and replisomes. To assess this, regions of interest (ROIs) were drawn around each SSB and DnaQ feature. Features were deemed to be colocalized if any portion of the ROI areas overlapped. Prior to irradiation, half of the SSB features did not overlap with a DnaQ focus (**Figure 3C; Supplementary figure 8**), indicating that many SSB features do not include an active replisome. We had previously observed that immediately after irradiation the number DnaQ foci decreased (**Figure 3B**) and the area occupied by SSB features significantly increased (**Figure 2C**). We observed that SSB features were even less likely to colocalize with a DnaQ focus at this timepoint; only 40% of SSB features overlapped with DnaQ (**Figure 3C**). Overlap frequencies gradually recovered to pre-irradiation levels over the next 90 min.

For those SSB and DnaQ ROIs identified as colocalized, we calculated both the mean percentage of overlapping area and the pair-wise centroid-to-centroid distance for each colocalized ROI pair, as a function of time (**Figure 3D–E**). Across all time points SSB foci consistently had 45–55% of their area overlapping with a DnaQ focus (**Figure 3D**), while SSB clusters decreased from 37% overlap with DnaQ (prior to irradiation) to 25% overlap (90 min after irradiation). Centroid-to-centroid distances between SSB clusters and DnaQ foci also became larger as a function of time (**Figure 3E**). Collectively these measurements support our initial inclination that DnaQ foci tend to localize at the outer edge of SSB features.

Analyzing colocalization from the perspective of DnaQ foci produced similar results. Prior to irradiation, 72% of DnaQ foci overlapped with an SSB feature (**Figure 3F**). Immediately after irradiation this increased to 80%. Taken together with the number of DnaQ foci (**Figure 3B**) and the SSB-centric colocalization results (**Figure 3C**), this is consistent with loss of some DnaQ foci and the remaining foci almost always being associated with an SSB feature. Overlap area was steady at 57-65% across the measurement (**Figure 3G**). Centroid-to-centroid distances between DnaQ foci and SSB features became larger as a function of time (**Figure 3H**), consistent with DnaQ localizing at the edge of SSB features.

### Formation of SSB clusters is largely independent of *recB*

The merging-splitting behavior presented in Figure 1, as well as the continued growth of SSB clusters observed in Figures 2–3, were suggestive of recombination events. To determine whether RecBCD-mediated homologous recombination played a role in the SSB-mTur2 dynamics we had observed, we next examined a single-color *ssb-mtur2* strain that lacked the *recB* gene. The *recBCD* recombination pathway is the main pathway involved in the repair of potentially lethal double strand breaks in *E. coli* [16]. The RecBCD protein complex binds to the end of a double-strand break before moving inward, unwinding, and resecting the DNA. Recognition of a chi sequence selectively suppresses degradation of the 3′ end. Thus, as RecB helicase continues to unzip and resect the 5’ strand, a ssDNA segment (chi tail), to which RecA can eventually be loaded, is formed. We hypothesized that SSB might also bind to the long tails created by RecBCD and thus cells lacking *recB* would produce fewer and smaller SSB features in microscopy images. For comparison we also analyzed a single-color *ssb-mtur2* strain in which the *recB* gene was left intact (*recB*^+^).

As before, cells were imaged in custom flow cells, with a continuous supply of aerated media, both before and after UV irradiation (fluence = 10 J/m^2^) *in situ* (**Figure 4A**). We monitored cell lengths (**Figure 4B**), intensities of cellular SSB-mTur2 signals (**Figure 4C**), as well as the number of SSB features per cell (**Figure 4D–E**). Prior to UV irradiation, similar morphology and SSB binding phenotypes were observed for *recB*^+^ and Δ*recB* strains. For both strains, cells measured roughly 5 µm in length (**Figure 4B**). The proportion of cells containing no DNA-bound SSB features was slightly higher for Δ*recB* cells than for *recB*^+^ cells (**Figure 4D**). The *recB*^+^ and Δ*recB* cells each contained two SSB-mTur2 features per cell on average. For both, the vast majority of features were foci (**Figure 4E**).

**Figure 4.**
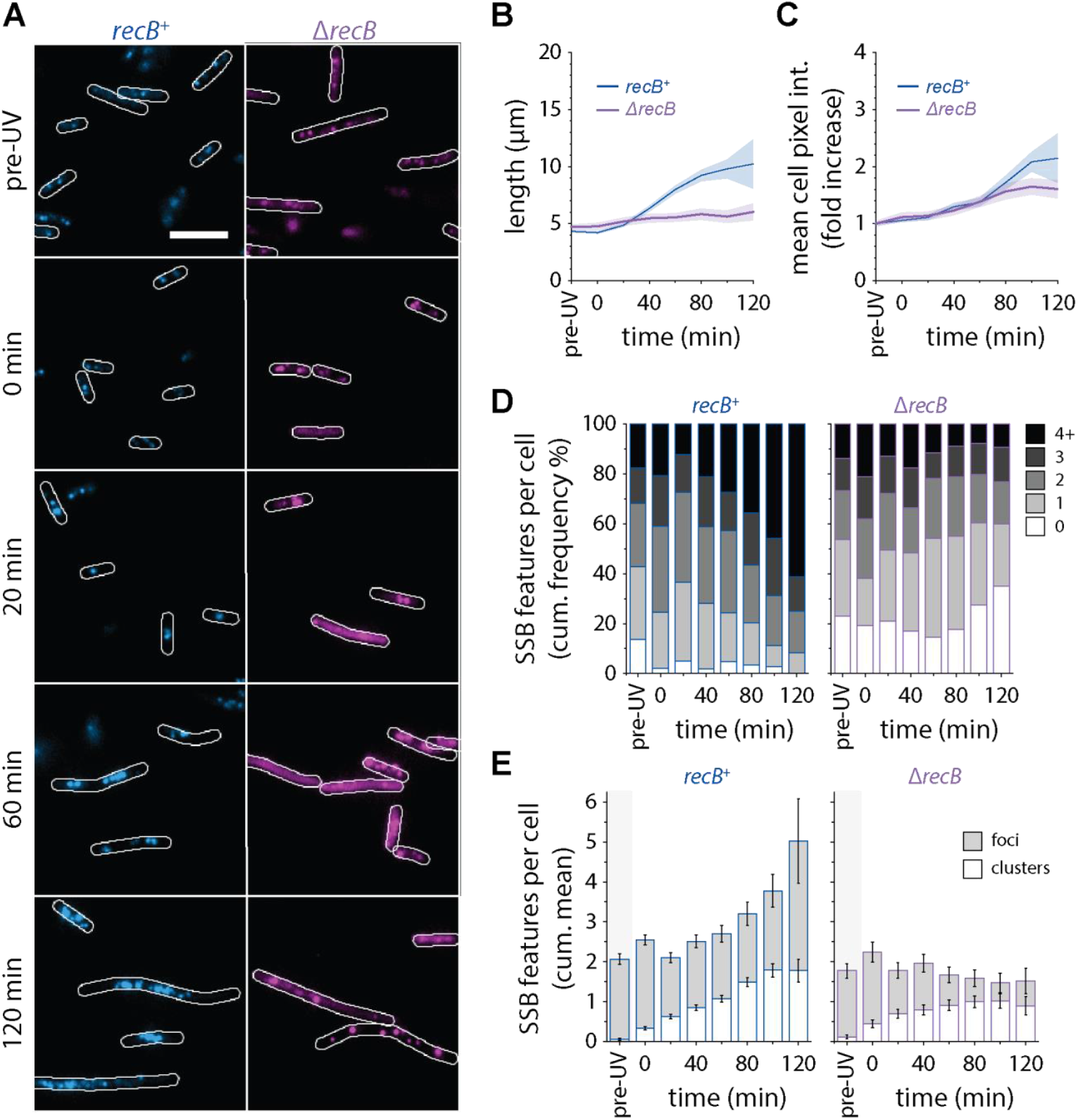
Effect of *recB* deletion on SSB-mTur2 binding phenotypes. **(A)** Time-lapse imaging of SSB-mTur2 in *recB*+ (blue, EAW1169) and Δ*recB* (pink, EAW1219) strains. Images shown are representative of those acquired at the indicated time post-UV exposure (fluence = 10 J/m2). Cells were manually outlined (white) using brightfield images. Scale bars indicate 3 µm. (**B-C**) Line plots of cellular length (B) and mean pixel intensity (C) over time. In all line plots, means associated with *recB*+ and Δ*recB* strains are indicated in blue and purple, respectively. Shaded regions indicate 95% confidence intervals. (**D-E**) Comparison of SSB-mTur2 binding over time in *recB*+ and Δ*recB* strains. (**D**) Population distribution of SSB cellular features (foci and clusters). (**E**) Quantification of bound SSB cellular features over time. The distribution of clusters (white) and foci (gray) are shown for each time point as a proportion of the total number of features. Error bars indicate 95% confidence intervals. (n ≥ 36 cells, N = 3)

Some clear differences between wild type and r*ecB*-deficient strains emerged following UV irradiation. Overall, cells lacking *recB* did not form extensive filaments in response to UV irradiation (**Figure 4B**), although a few elongated cells were observed (**Figure 4A**). Increases in the mean SSB pixel intensities of Δ*recB* cells were similar to those observed in the *recB*+ strain (**Figure 4C**). UV irradiation stimulated the formation of SSB features in the *recB*^+^ cells (**Figure 4D**). Immediately after exposure to UV, the proportion of cells without detectable DNA-bound SSB features dropped from 14% to 2% and remained low 2 h after irradiation (**Figure 4D**, left panel). No such drop was observed for Δ*recB* cells (**Figure 4D**, right panel). Instead, a growing population of featureless cells (containing only diffuse SSB signal) was observed in the 2 h following irradiation (**Figure 4A, D**). Except for a slight increase immediately following irradiation, the total number of SSB features present in Δ*recB* cells did not differ significantly from pre-UV values (**Figure 4E**, right panel), in contrast with steady increases observed for *recB*^+^ cells (**Figure 4E**, left panel). On the other hand, the proportion of SSB clusters in Δ*recB* cells increased steadily in response to UV (**Figure 4E**, right panel), as was observed for *recB*^+^ cells (**Figure 4E**, left panel). Thus, despite some notable differences in the behaviors of *recB*^+^ and Δ*recB* cells, the formation of SSB clusters appears to be independent of *recB*.

## DISCUSSION

In this study we present six main conclusions with respect to visually detectable SSB features in *Escherichia coli* growing in relatively rich media: 1) Under normal growth conditions a small number of SSB foci (~8%) are conspicuously brighter than others (**Figures 1–2**). 2) In cells exposed to a flash of UV light, a larger proportion (~17% %) of SSB foci are bright and additional SSB clusters form (**Figures 1–2**). A much higher fraction of cells produced one or more bright foci at some point during a replication cycle. 3) Bright SSB foci are highly dynamic on the low-minutes timescale, forming and dissipating *via* either a merging–splitting pathway or a brightening–dulling pathway (**Figure 1A**). 4) SSB features become larger with increasing recovery time after UV irradiation (**Figure 2**). 5) Replication fork markers co-localize with many but not all SSB foci and tend to localize at the outer edge of large SSB features (**Figure 3**). 6) The formation of SSB clusters is largely independent of the *recB* pathway (**Figure 4**). The many features that do not co-localize with replisomes, particularly after UV irradiation, indicate that intermediates associated with processes such as the repair of mismatches, double strand breaks, or post-replication gaps are both common and quite readily detected by this approach.

The results of this study provide a first look at multiple types of SSB features being formed in *E. coli* cells under normal growth conditions, and in cells exposed to UV light. The study as designed provides an important baseline for an approach that has the potential to reveal much about important aspects of DNA metabolism. The method described here also provides an important complement to genomic examination of ssDNA locations via nondenaturing bisulfite treatment followed by deep sequencing [17]. That previous work [17] has established that most readily detectable ssDNA gaps are associated with replication. In addition, the overall amount of ssDNA present at a given time in log phase cells (~1.3% of the total DNA in log phase cells or ~60,000 nucleotides) is significantly greater than expected for generation of ssDNA at the replication forks. The excess of SSB features observed away from replisomes in our study correlates well to this observation. Additionally, the levels of ssDNA increased about three-fold following UV irradiation, a result that again correlates well to the Pham *et al*. study [17]. Thus, we can now see features of the ssDNA genomic landscape by two complimentary methods, features that were previously largely inaccessible.

Although the identities of the various types and patterns of foci formation cannot be definitively assigned at this stage, many of the features exhibit characteristics that are highly suggestive of various ssDNA-containing DNA repair intermediates. Possible identities for the different types of SSB features are discussed below but a summary of key results at the outset is useful. There are more SSB features than replisome foci, a situation most evident after UV irradiation. A significant fraction of the SSB features can thus likely be attributed to repair of mismatches, double strand breaks, or post-replication gaps. The strain lacking *recB* function provides an example of how this approach can be utilized to catalogue and categorize DNA metabolism events. Similarities in SSB features for WT and Δ*recB* cells prior to UV irradiation suggests that relatively few incidents of double strand break repair occur under our normal growth conditions. After UV irradiation, an increase in total SSB foci can be attributed to double strand break repair, as the increase is not seen in a *recB* deletion. However, an observed increase in the number of SSB clusters occurs independently of RecB. The additional observation that centroid-centroid distances between replisomes and SSB clusters increase over time after irradiation suggests that these SSB clusters expand away from the replisome. We very tentatively propose that at least some of these larger SSB clusters reflect the formation of post-replication gaps.

### Formation and dissipation of bright SSB foci on the low-minutes timescale

In the absence of exogenous damage (i.e. prior to UV irradiation) >90% of the SSB features we observe in the current study are relatively dull foci (**Figure 1A, Figure 2A**). Half of these foci colocalize with a marker for DNA polymerase III (**Figure 3C**). The most obvious explanation is that these replisome-proximal features represent SSB bound to the template lagging strand within replication forks. However, it is important to note that light microscopy has inherent resolution limitations and that some of these foci might also arise from SSB molecules bound at post-replication gaps located close behind the replisome.

The identity of the remaining 50% of SSB foci that do not colocalize with DNA polymerase III (**Figure 3C**) is less clear. One possibility is that these foci originate from replication forks that have collapsed, leaving the SSB behind. Replication fork collapse appears to be relatively common in bacteria; measurements in *E. coli* and *Bacillus subtilis* cells indicate that >40% of replication forks are disassembled at any given time due to conflicts with transcription complexes [59].

In the current study we observed that under normal growth conditions there is a sub-population of foci (~8%) that are brighter than the rest. This suggests that ssDNA species that are much longer than Okazaki fragments are produced relatively frequently. Under these conditions, the most common ssDNA species in cells that are long enough to reliably attract SSB binding are likely to be post-replication gaps and ssDNA intermediates produced during double strand break or mismatch repair. Post-replication gaps and mismatch repair gaps are likely to be long enough to accommodate multiple tetramers of SSB (15-30 tetramers in the 65 mode for a gap in the range of 1-2 kilobases). Post-replication gaps formed at the lagging strand would be expected to have a similar distribution of lengths as the Okazaki fragments produced during DNA synthesis. In vitro measurements suggest that at their largest, mismatch-repair gaps are comparable in length to Okazaki fragments [60]. If this is true *in vivo* under our growth conditions, then mismatch repair gaps are unlikely to be the source of the bright SSB foci.

There is no information available to estimate the size of post-replication gaps. The results of two recent studies [61, 62] indicate that the RecJ nuclease [63] acts within the early stages of post-replication gap repair. If RecJ were to expand post-replication gaps by degrading the 5′ gap end of the dsDNA downstream, this would create a binding site for large amounts of SSB. The bright SSB foci we observe (**Figure 1**) may therefore be consistent with gap-expansion products formed during the repair of post-replication gaps, especially those bright foci produced *via* brightening (as opposed to merging). The brightening process takes place on the low-minutes timescale (**Figure 1A**). These dynamics may also be consistent with gap expansion by RecJ. Within 1–2 min the RecJ nuclease can degrade 1–2 knt of ssDNA [64], conceivably providing space for an additional 15–30 SSB tetramers to bind (in the 65-mode). This would in principle be sufficient to drive the brightening process that we observe. Finally, one might expect that DNA lesions introduced by exposing cells to UV light could trigger the formation of additional post-replication gaps. Indeed, we observe a rapid increase in the number of bright SSB features following exposure to UV light (**Figure 1D–E**). Upon UV irradiation, the total number of SSB foci increases, something that does not occur when *recB* is deleted. Much of the increase can thus be attributed to double strand gap repair as may occur when replisomes encounter template strand discontinuities associated with ongoing nucleotide excision repair of UV damage. However, the number of bright foci seen after UV irradiation increases in a manner that is independent of RecB, further suggesting that many and perhaps most of these are associated with post-replication gaps.

A key step in the repair of post-replication gaps is loading of the RecA recombinase onto the SSB-coated ssDNA in a reaction mediated by RecO in complex with RecR [56, 65]. Replacement of some or all of the SSB by RecA in this manner would be consistent with the dulling of bright SSB foci that is observed in the current study (**Figure 1A**). In the same data we also observed SSB foci merging and splitting. One possibility is that this is driven by homologous recombination. Perhaps regions of chromosomal DNA (and their associated SSB features) that are normally spatially distinct are brought into close proximity during recombinational repair, leading to the merging of SSB foci. Following repair the DNAs separate from each other, leading to focus splitting. A similar set of events have been previously observed during recombinational repair of double-strand breaks [66, 67]. A follow-up study exploring the genetic dependencies of the brightening, dulling, merging, and splitting behaviors is currently underway.

### SSB clusters formed in response to UV light

In addition to regular foci and bright foci, we observed large SSB features that appeared to be comprised of multiple partially overlapping foci. These features became larger during the 120 min recovery period. A series of complex phenomena are expected to occur in the period following UV irradiation, and many of these could produce substrates for SSB. One possibility is that the large features represent multiple gap-repair intermediates that become aggregated together. A follow-up study exploring the genetics of large SSB features is currently underway.

Our work demonstrates a viable experimental approach to the real-time observation of the formation and processing of single-stranded DNA gaps in individual bacterial cells. The methods reported here will provide a foundation for future work understanding the molecular mechanisms underlying gap creation and processing in bacteria systems.

## AVAILABILITY

*Single-Molecule Biophysics* is a Fiji plug-in available in the GitHub repository (https://github.com/SingleMolecule/smb-plugins). *Process Raw Images, ChangePoint Peaks, MeasureROIsAllFrames*, and *Particle Analysis Suite v4* are custom Fiji macros available in the GitHub repository (https://github.com/chaireontop/MC-Custom-Macros)

## ACKNOWLEDGEMENTS

The authors thank Myron Goodman and Roger Woodgate for helpful advice and feedback.

