## Supplementary Material for "Spatiotemporal Dynamics of Single-stranded DNA Intermediates in *Escherichia coli*"

SUPPLEMENTARY FIGURES

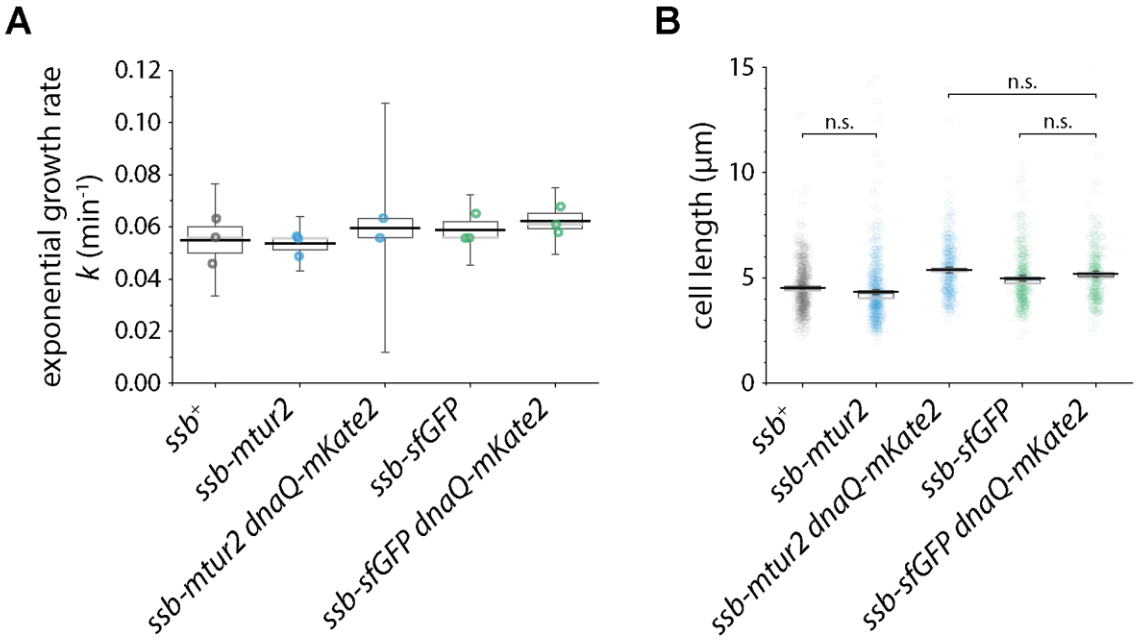


**Supplementary figure 1. Growth and morphology of cells expressing SSB fusions.** (**A**) Growth rates of SSB-IDL chromosomal fusion strains. The growth rates of single- and dual-color *ssb-mtur2* and *ssb-gfp* strains did not differ significantly from wild-type MG1655 or each other (One-way ANOVA, p-value = 0.4872). OD600 readings were measured in a 96-well plate in technical triplicate (see Methods for more details) and fit to a Richard's curve to extract growth parameters. Box plots indicate 95% CI (whiskers), SE of the mean (box), median (gray line), and mean (black line) for each quantity. (**B**) Box plots of cell lengths for cells expressing unlabelled ssb (black), ssb- mtur2 (blue), and ssb-gfp (green) containing strains. Plots indicate mean (black line), median (gray line), 95% CI (whiskers) and SE of the mean (box). A Tukey test was used to determine significance to $p < 0.05$.

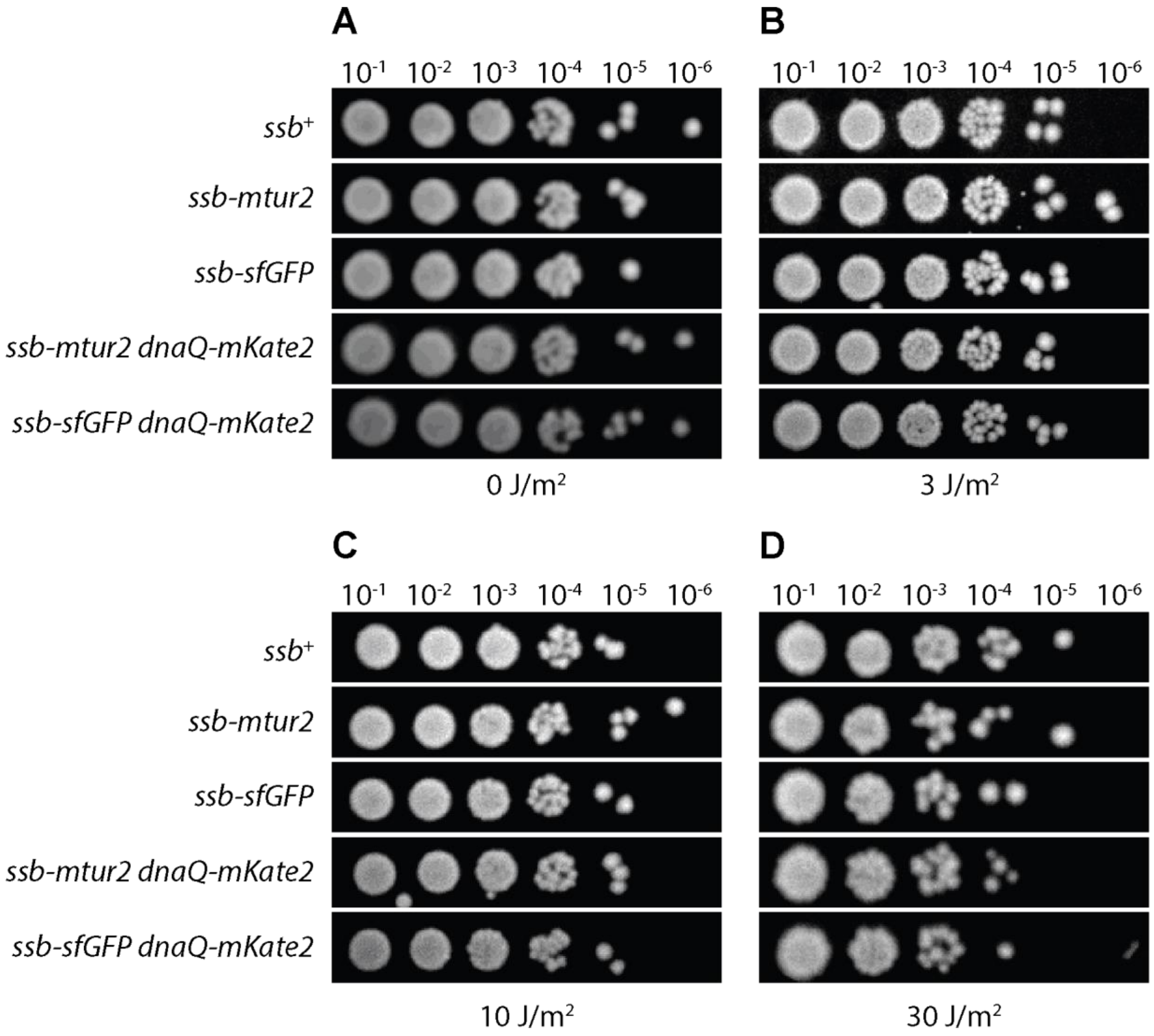


**Supplementary figure 2. Representative images of UV survival spot plates.** Cells expressing SSB-mTur2 or SSB-GFP demonstrate no additional UV sensitivity than cells expressing wild type SSB. Overnight cultures were grown in LB medium, diluted 100-fold and allowed to grow for 120 min. Cells were irradiated with doses of (**A**) 0, (**B**) 3, (**C**) 10, or (**D**) 30 J/m2 with 254-nm light. These samples were serial diluted and plated onto LB agar plates. Experiments were minimally conducted in biological duplicate.

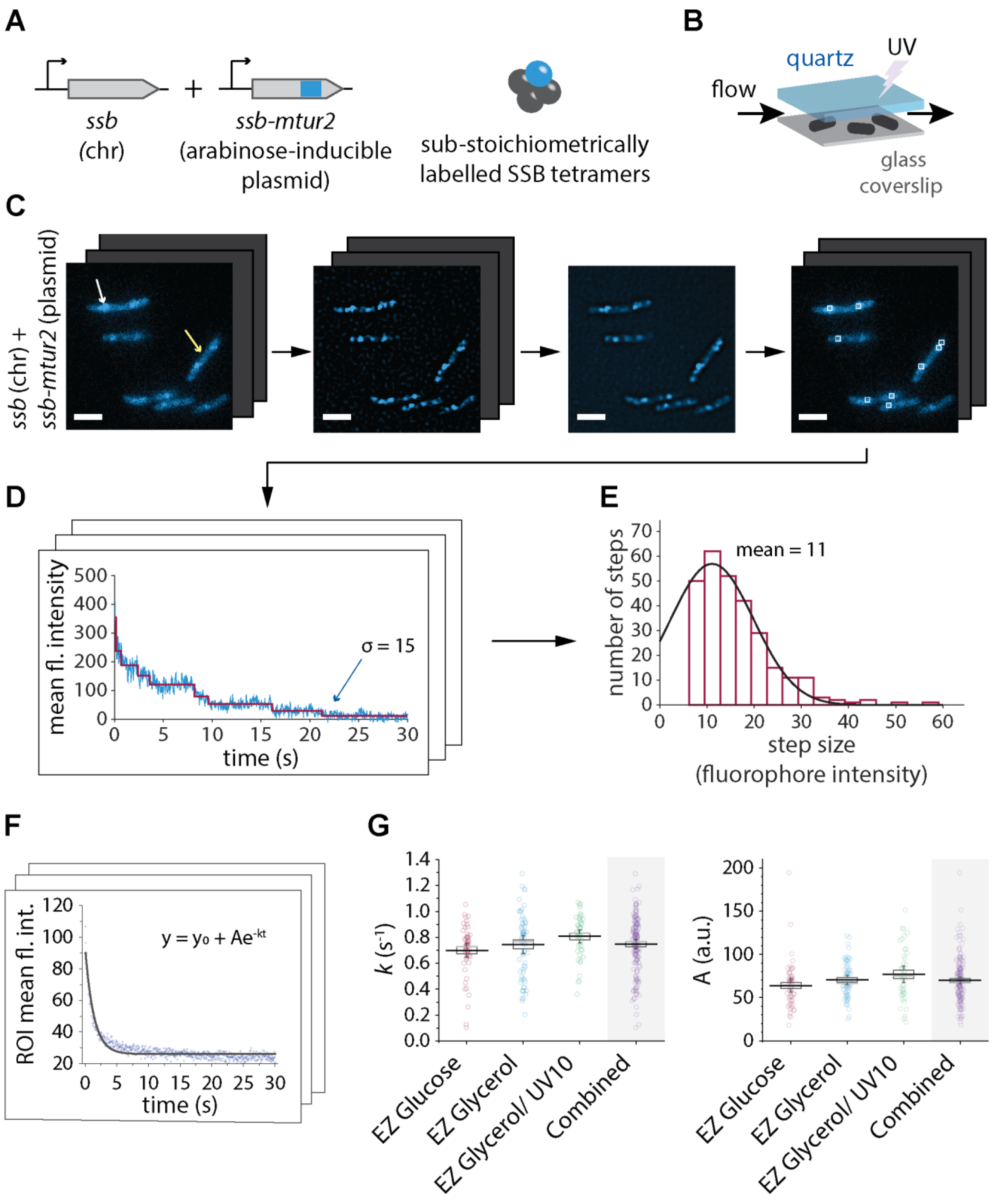


**Supplementary figure 3. Calibration measurements for calculation of intracellular SSB concentrations.** (**A–E**) Measuring the mean intensity of a single mTur2 molecule. (**A**) The *ssb-mtur2* internal fusion was expressed under an arabinose-inducible promoter from a pBAD vector (pEAW1197) in wild-type MG1655 cells. At 0.002% L-arabinose, it is expected that SSB tetramers are a mixture of dark chromosomally expressed SSB (gray) and fluorescent plasmid expressed SSB-mTur2 (cyan). (**B**) Cells expressing SSB-mTur2 from an arabinose-inducible plasmid were grown to early exponential phase with an induction period of 30 min (0.002% arabinose) and loaded into a flow cell. Cells were UV irradiated (fluence 10 $J/m^2$) to promote SSB binding, then imaged under constant supply of aerated growth medium with illumination from a 458 nm laser. (**C**) Fluorescence image stacks of SSB-mTur2 (cyan) captured both long-lived protein binding (white arrow) and diffusive transient movement (yellow arrow). To isolate longer-lived DNA bound foci, image stacks were first corrected for excitation inhomogeneity (left) then discoidal filtered (middle left). The average projection was then taken across all frames (middle right). Mean intensities were then measured at the identified peak locations in the original flattened stack (right). Scale bars indicate 3 μm. (**D**) A kinetic change point algorithm [1] was used to distinguish discrete molecule steps (red) from noise in each mean intensity photobleaching trajectory (blue). Only final photobleaching steps (blue arrow) were used in the calculation of the mTur2 single-molecule mean pixel intensity. (**E**) Histogram of mTur2 bleaching trajectory final step sizes (Istep) (red, n = 281 trajectories). An mTur2 mean pixel intensity of 11 arbitrary units (95% CI: 10 – 13) was derived from the single-term Gaussian (black line) fit to the binned data. (**F–G**) Quantitation of cellular autofluorescence. (**F**) Fluorescence image stacks of label-free wild-type cells were used to quantify cellular autofluorescence. The mean pixel intensity of each cell was plotted over time (blue) and fit to a single-exponential photobleaching curve (dark gray) with photobleaching rate *k*, offset $y_0$ and amplitude *A*. For each cell, *A* was taken as a measure of the mean autofluorescence pixel intensity. (**G**) Box plots of relevant photobleaching parameters for MG1655 cells grown in EZ rich defined media supplemented with 0.2% glucose (EZ Glucose) (red), EZ rich defined media supplemented with 0.2% glycerol (EZ Glycerol) (blue), and EZ Glycerol with UV irradiation (fluence 10 $J/m^2$, green) ($n \geq 44$ cells). Box plots indicate 95% CI (whiskers), SE of the mean (box), median (gray line), and mean (black line) for each quantity. As *k* and *A* were not statistically different across the various growth conditions (One-way ANOVA, p-value (k) = 0.08208, p-value (A) = 0.0543), the mean value for the combined set was used when calculating intracellular SSB concentrations.

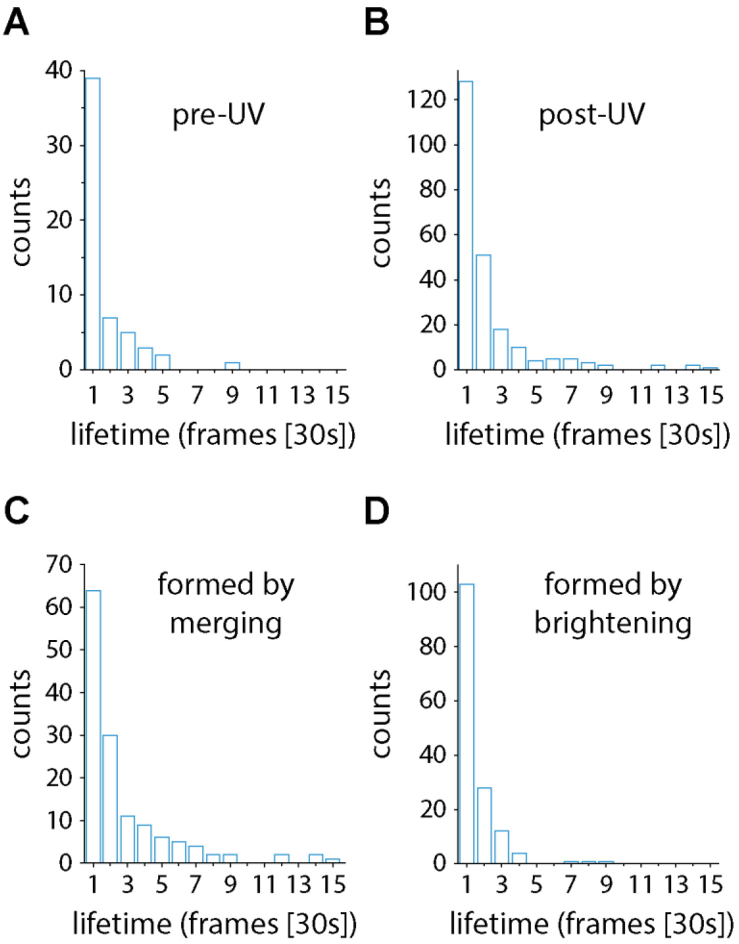


**Supplementary figure 4. Lifetimes of bright SSB foci.** (**A–D**) Histograms of time spent in the bright state for SSB-mTur2 foci. Bin labels indicate the number of frames for which bright foci were detected. The interval between frames was 30 s. (**A**) Bright foci observed prior to UV irradiation. (**B**) Bright foci observed after UV irradiation. (**C**) Bright foci that formed by the merging of multiple foci. (**D**) Bright foci that formed by brightening of weaker foci.

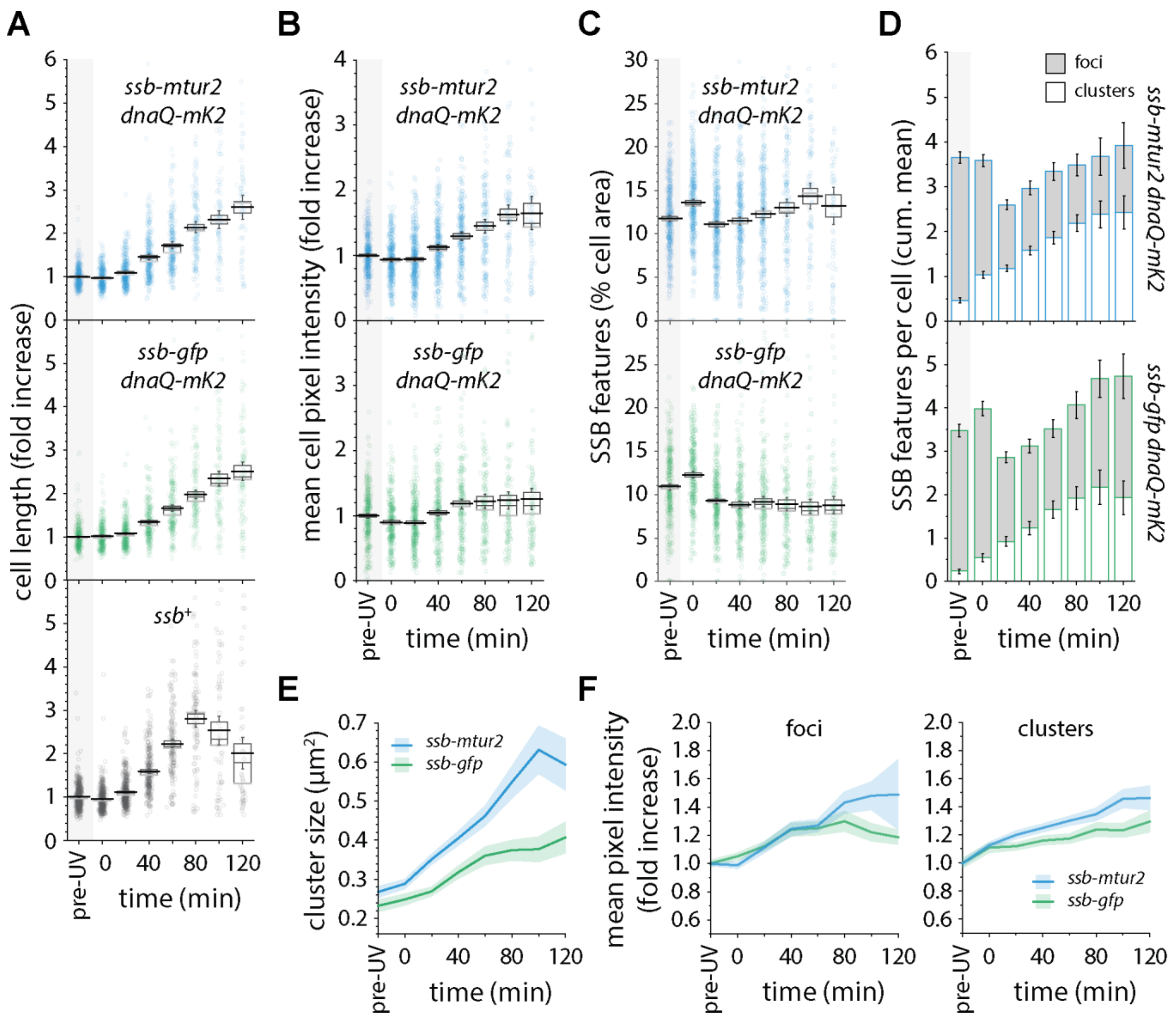


**Supplementary figure 5. Quantification of cell length, cellular mean pixel intensity and DNA-bound SSB features over time in response to UV irradiation.** Box plots of the measured cell length (**A**), cellular mean pixel intensity (**B**), and proportion cell area occupied by bound features (**C**) over time. Plots indicate the mean (black line), median (gray line), 95% CI (whiskers), and SE of the mean (box). Data for *ssb-mtur2* (MEC189), *ssb-gfp* (MEC191), and *ssb*$^{+}$ (MG1655) containing strains are indicated in blue, green, and black, respectively. (**D**) Quantification of bound SSB cellular features over time. The distribution of particles (white) and peaks (gray) are shown for each time point as a proportion of the total number of SSB features per cell. Error bars indicate 95% confidence intervals. (**E**) Particle size over time. Means indicate the average particle ROI size at the indicated time after UV irradiation. (**F**) Mean pixel intensities of DNA bound SSB features over time for peaks (left) and particles (right). Means indicate the average pixel intensity of peaks and particles as fold increases from the pre-UV intensity. In all line plots, means associated with ssb-mtur2 and ssb-gfp containing strains are indicated in blue and green, respectively. Shaded regions indicate 95% confidence interval regions. ($n \geq 46$ cells, 47 particles, and 102 peaks per time point; $N \geq 2$).

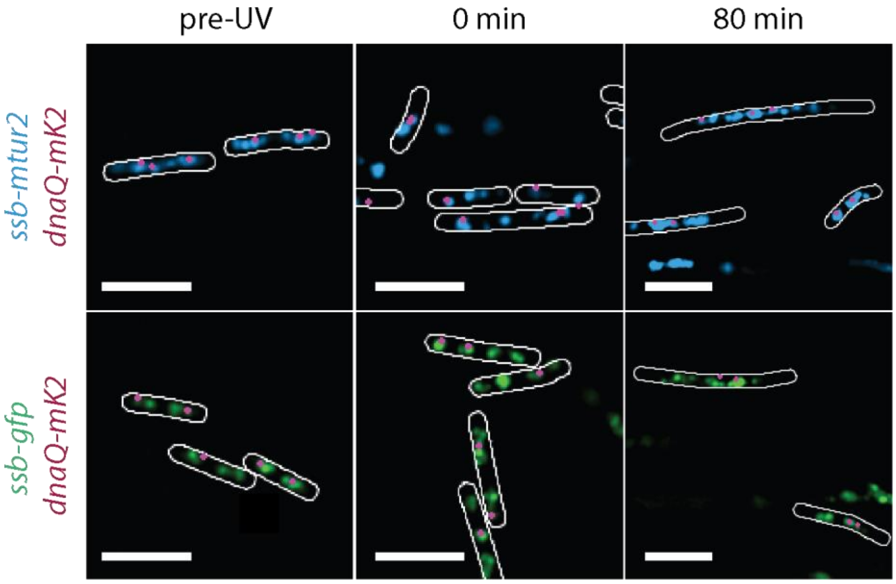


**Supplementary figure 6. Localization of DnaQ-mKate2 with respect to SSB-mTur2 and SSB-GFP.** Time-lapse images of *ssb-mtur2 dnaQ-mK2* (MEC189) and *ssb-gfp* dnaQ-mK2 (MEC191) cells were acquired before and after UV irradiation (10 J/m$^2$) *in situ*. Discoidal filtered images of SSB-mTur2 (top, blue) and SSB-GFP (bottom, green) are representative of those acquired at the indicated times after exposure. Cell outlines (white) were manually drawn based on brightfield images. Locations of DNA-bound replisome foci are indicated by circular spots (magenta). Scale bars indicate 5 μm.

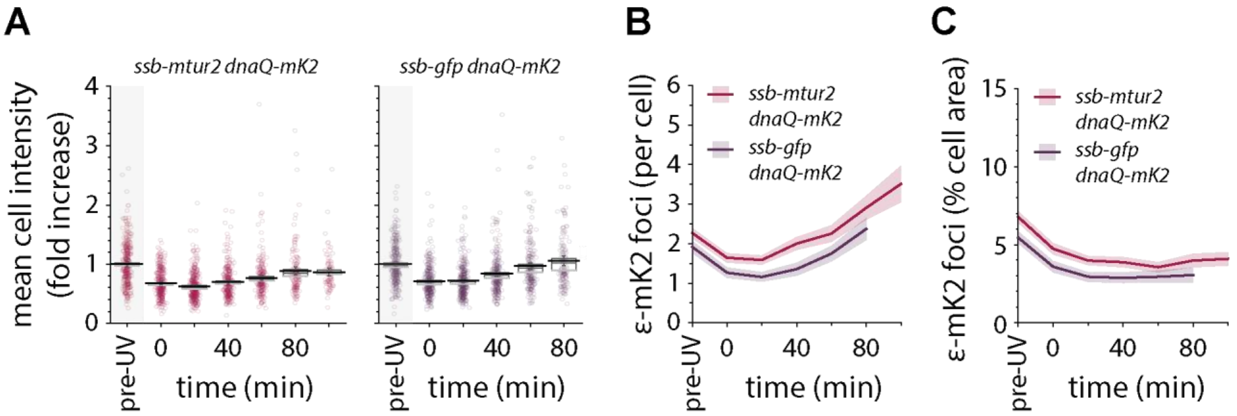


**Supplementary figure 7. Characterization of DnaQ-mKate2 signal over time.** (**A**) Box plots of the DnaQ-mKate2 cellular mean pixel intensity over time for *ssb-mtur2 dnaQ-mKate2* (MEC189, left panel) and *ssb-gfp dnaQ-mKate2* (MEC191, right panel) cells. Plots indicate the mean (black line), median (gray line), 95% confidence intervals (whiskers), and SE of the mean (box). (**B-C**) Quantification of DnaQ-mKate2 foci over time as foci per cell (**B**) and the proportional cell area (**C**). In all line plots, means associated with ε-mK2 activity in MEC189 and MEC191 strains are indicated in red and violet, respectively. Shaded regions indicate 95% confidence interval regions. (n ≥ 86 cells).

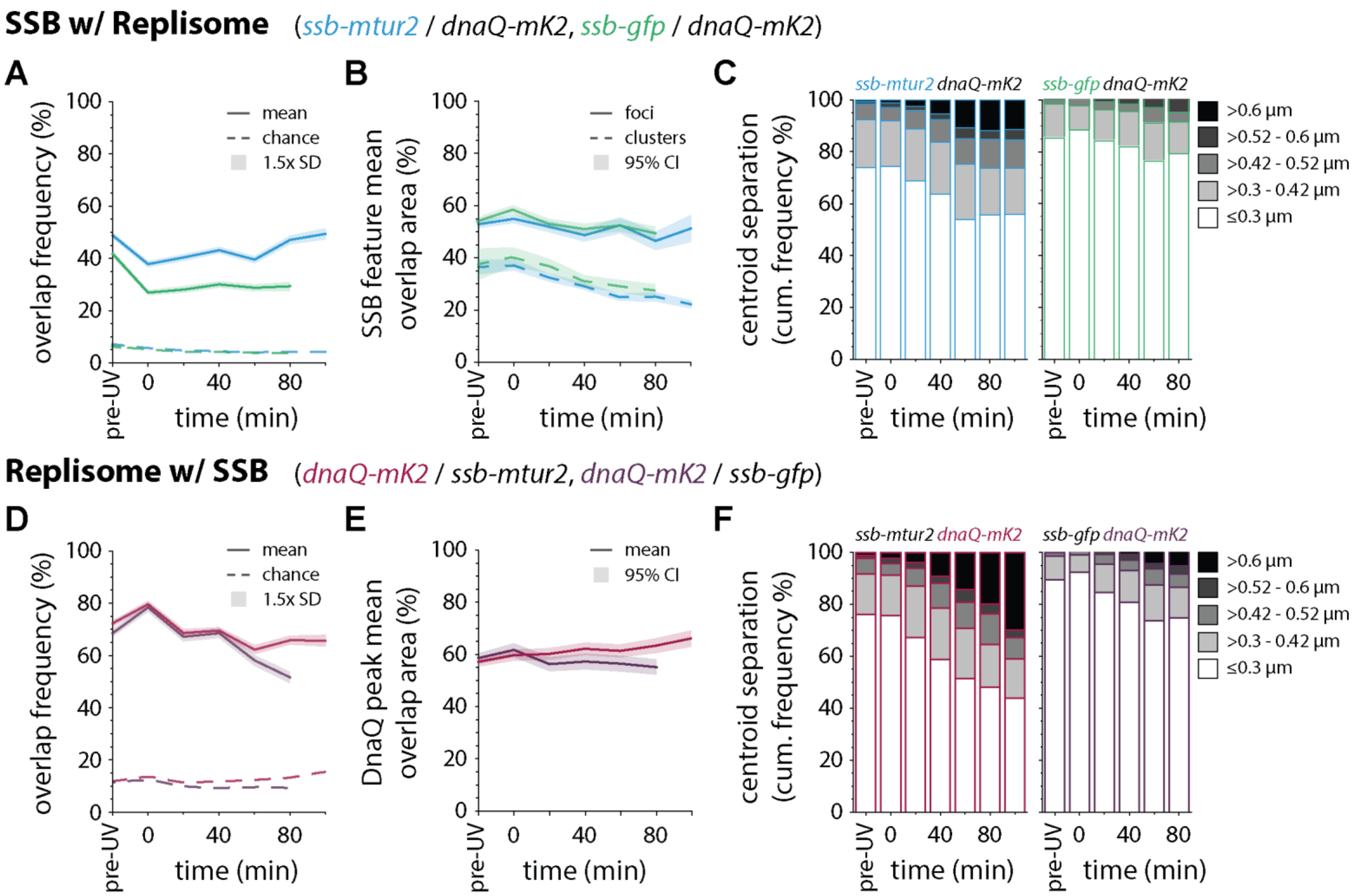


**Supplementary figure 8. Analysis of SSB/replisome spatial overlap.** In all plots values associated with *ssb-mtur2 dnaQ-mKate2* are presented in blue and red. Values associated with *ssb-gfp dnaQ-mKate2* are presented in green and violet. (**A, D**) Plots of overlap frequencies of SSB with replisomes (**A**) and replisomes with SSB (**D**). Chance colocalization in each case is indicated by a dashed line. Shaded regions indicate values within 1.5× SD of the mean. (**B, E**) Quantification of area overlap between colocalized features. The mean percentage of overlapping area for each region of interest (ROI) associated with colocalized SSB and replisome peaks (solid lines) and SSB particles (dashed lines) was plotted over time. Shaded regions indicate 95% confidence intervals. (**C, F**) Quantification of overlapping feature centroid - centroid separation. Radii ranges correspond to 0.09 µm$^2$ area shells.
